# Weak dispersal and landscape size inevitably promote local biodiversity in heterogeneous metacommunities of competing species

**DOI:** 10.64898/2026.02.16.706088

**Authors:** Frederik De Laender, Andrew Gonzalez, Olivia Bleeckx, Dieter Ebert, György Barabás

## Abstract

We provide a general theoretical explanation for a longstanding result in spatial ecology: weak dispersal among habitat patches promotes local biodiversity. Using analytical approximations of spatial Lotka–Volterra competition models, we show that species persistence in heterogeneous landscapes can be expressed as a function of regional abundance and local invasion growth rates. We further demonstrate that local multispecies coexistence is governed by the feasibility domain, linking spatial coexistence to a structural property of nonspatial competitive systems. Together, these results explain why weak dispersal increases local species richness and why this effect strengthens with landscape size.

We test these predictions using numerical simulations and find that the theory breaks down only when both dispersal and competitive interactions are very strong, in which case dispersal has a unimodal effect on coexistence. In contrast, landscape size retains a positive effect on coexistence whenever an effect is detectable. We then apply the theory to long-term data from a natural *Daphnia* metacommunity. We detect strong preemptive competition among species and find no detectable effect of dispersal rate on local coexistence, whereas species co-occurrence increases with local landscape size, as predicted by theory. Together, our results identify how dispersal, interaction strength, and landscape size jointly regulate biodiversity in competitive systems.

## 1 Introduction

The effect of dispersal—the movement of organisms from one location to another—on local species richness depends on a variety of factors. Spatial heterogeneity in environmental conditions [1, 2], the strength of local species interactions [3, 4], and the directionality [5], capacity [6], and density-dependence of dispersal rates [7], all have been reported as significant factors. However, despite this multitude of factors shaping the effect of dispersal on local species richness, one finding consistently emerges from most experimental [8, 9] and modeling studies [10–14], as well as from meta-analyses [15, 16]: in heterogeneous landscapes, weak dispersal tends to promote local species richness.

The verbal explanation for weak dispersal affecting local richness positively is that weak dispersal alleviates local species interactions, which thence increases local richness. Simulation models have played an important role in providing numerical evidence for this mechanism [11, 17]. One approach extends the classical Lotka–Volterra equations for *n* interacting species with dispersal [12, 18]. These models track the population density 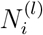 of every species *i* in every patch *l*, across a spatial network of *p* patches inhabited by *n* interacting species:

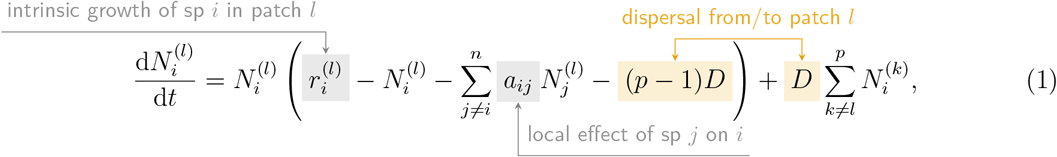

where 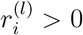 is the growth rate of species *i* in patch *l, a*_*ij*_ is the effect of species *j* on *i*, and *D* is the dispersal rate, which measures how quickly individuals move across patches.

An important drawback of Eq. 1 is that it is analytically intractable for diverse communities and thus hard to draw general insights from. Available analytical results for this model therefore focus on communities with few species [10, 19, 20], highlighting how spatial heterogeneity in competitive ranking can enable coexistence [10], and how different dispersal mechanisms can modify coexistence outcomes [7]. Other results of models based on Eq. 1 ignore the discrete nature of the spatial network by relying on a reaction-diffusion formalism [21, 22]. One exception is the contribution by Levin [3], showing that dispersal promotes invasion into a patch if the local conditions of that patch permit growth of the invader. How these results relate to long-term persistence has remained unclear, however. More parsimonious and therefore analytically tractable alternatives to Eq. 1 include patch occupancy community models [23–26], where one tracks the fraction of patches that species occupy in a landscape. However, this research program cannot explain why weak dispersal has a positive effect on local richness, because its focus is on persistence at the landscape level, not on local abundances [2]. In summary, to date, there is no general explanation of the positive effect of weak dispersal on local richness in large communities.

Here we present new analyses that explain the well-documented positive effect of weak dispersal on local species persistence in heterogeneous landscapes. Considering dispersal to be weak and species growth rates to vary across the landscape, we are able to analyze Eq. 1. We show that when dispersal is sufficiently weak to have a linear effect on local population density, quantifying the roles of dispersal, landscape size, and local species interactions is within reach. We first recover the result that weak dispersal inevitably promotes persistence (measured by the fraction of patches in which a randomly selected species has positive density). This effect is stronger in landscapes with more patches and weaker interactions. We then show the implications of these results for coexistence, quantified via the fraction of patches in which all *n* regionally available species coexist. We show how this multispecies extension of the patch occupancy concept, initially developed for metapopulations [24, 26–28], relates to feasibility, a classic quantity in contemporary coexistence theory [29, 30]. Further, we show that persistence and patch occupancy are analytically predictable when the mean growth rate across all patches is the same for all species (i.e., species are regionally equivalent), and all interspecific interactions are exactly equal (diffuse species interactions, [31, 32]). Model simulations show that this prediction is exact, and that our conclusions continue to qualitatively hold when relaxing our analyses’ underlying assumptions, unless interspecific interactions are strong and dispersal is so large as to have a nonlinear effect on population densities. Finally, we compare our findings with a metacommunity of *Daphnia* species, sampled across hundreds of rockpools on the Tvärminne archipelago (Finland). This analysis suggests strong interspecific interactions and shows that species co-occurrence is more likely in pools surrounded by a greater number of pools.

## 2 Methods

### 2.1 Analyses

Moving from observing to understanding the roles of dispersal and species interactions is challenging because mathematically analyzing Eq. 1 is not trivial. To solve this problem, we consider dispersal to be weak. Specifically, when 0 ≤ *D* ≪ 1, the equilibrium density of species *i* in some patch *l*, 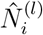, is approximated by a linear function. It is the sum of what it would have been without dispersal (which we call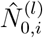, where the 0 subscript represents absence of dispersal, used throughout the manuscript) and the amount of density contributed per unit of dispersal (which we call 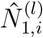), leading to 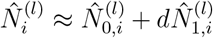, where *d* equals *D* after rescaling. This rescaling is needed to remove units and permit the small dispersal approximation. Doing so leads to an implicit equation for 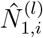 (Supporting Information, Section 1.1), which we analyze to study how dispersal affects local species persistence, defined as the fraction of patches in which the given focal species *i* persists. This fraction is equivalent to the probability that a random species persists in a random patch (i.e., the probability that 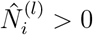).

We then calculate the probability that 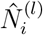 is larger than some threshold, given spatial variation of the intrinsic growth rates 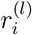. To obtain an explicit equation for this probability, we introduce the following simplifying assumptions. First, all the 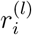 are i.i.d. random variables following a standard normal distribution, truncated to the positive real numbers, and standardized such that the mean across species within any given patch, and its mean across patches for any given species, is 1. This introduces spatial heterogeneity because species grow differently in different patches (as in [33]), but also implies that species are regionally equivalent: the mean growth rate across all patches will be the same for all species. Second, we assume a diffuse and weak interaction scheme: all pairwise species interactions are exactly equal (*a*_*ij*_ = *a* for all *i* ≠ *j*) and weaker than intraspecific interactions (*a < a*_*ii*_ for all *i*). Persistence probabilities were calculated using the methods specified in Supporting Information, Section 1.2.3, for a regional pool of *n* = 10 species.

We also compute community-level patch occupancy as the fraction of patches in which all *n* = 10 coexist. This fraction is equivalent to the probability that all species persists in a random patch. To this end, we use the methods from Supporting Information, Section 1.2.3 and [34].

### 2.2 Simulations

We ran simulations with Eq. 1 to test how relaxing the assumptions of our analyses would change the effect of dispersal on local species richness. We adopted a fully-factorial simulation design by varying *p* and *d* (values shown in Fig. 2), the across-pairs mean *a* (= 0.2, 0.5, 1) (i.e. going from weak to preemptive competition [20]) and variance of interspecific interaction strength (coefficients of variation of 0 or 0.2), whether these interactions varied across patches (yes or no), whether there was regional equivalence of the growth rate (yes or no), and if dispersal happened between all patch pairs (well-mixed scenario) or obeyed exponentially declining dispersal, whereby nearby patches exchange more individuals than farther ones. We ran 100 replicate simulations per factorial combination and fixed the number of regionally available species at *n* = 10.

To vary interactions across patches, we multiplied each interaction with a coefficient randomly sampled from a uniform distribution with minimum 0.8 and maximum 1.2. To break regional equivalence of the growth rate, we multiplied a randomly-picked species’ regional mean growth rate by 1.5, and divided that of the other species by that same amount. Making dispersal follow an exponential dispersal kernel involved randomly generating coordinates for every patch and characteristic distances for every species, and based on these calculating dispersal rates between two patches as *e*^−*s/c*^, where *s* is the distance between the two sites and *c* is the characteristic distance the species can cover in one unit of time. We then standardized all dispersal rates to have a mean equal to the value for the well-mixed scenario.

For every factorial combination and replicate, we sample *n* strictly positive growth rates for every patch (by sampling *p* times *n* growth rates from a standard normal distribution), take the absolute values of these rates, and iteratively normalize such that the mean across species within any given patch, and its mean across patches for any given species, were both 1. We seeded species at a density numerically equivalent to one tenth of their local growth rate, and simulated the dynamics of the metacommunity until reaching equilibrium using the R [35] package `rootSolve` [36, 37]. We considered a species extinct when its final density fell below the threshold of 10^−3^. We then calculated patch occupancy as the fraction of patches in which all *n* species persisted.

To facilitate comparison with the analytical results and be able to tell what levels of *d* can be considered sufficiently weak, we ran an additional simulation to evaluate within which range *d* has a linear effect on local density. These results are in Supporting Information, Fig. S1.

### 2.3 *Daphnia* data analysis

The occurrence of three *Daphnia* species (*D. magna, D. pulex, D. longispina*) in freshwater rock pools on the skerry islands of the Tvärminne archipelago (Finland) has been censused twice per year since 1982 [38–40]. These islands form seven spatial clusters, with rock pools connected by wind-driven dispersal of resting eggs following pool desiccation [41]. We attributed islands to a cluster based on proximity: islands closer than 100 meters to each other were considered as a single cluster. We assume dispersal between two island clusters is negligible. We use simulated desiccation rates for every pool and year from Altermatt et al. [42], as a proxy of the dispersal rate. Within each cluster, we first measured the distance between each pair of pools. We then counted, for each pool, the number of neighbor pools within a given distance. We carried out our statistical analysis for distances of 50 and 100 meters to test the robustness of the obtained results. Co-occurrence of all three species was very rare, having been observed in only 0.33 % of all sampling events. We therefore focus on co-occurrence of species pairs in what follows, and only consider pools with either one or two species.

Given their important role for local coexistence in metacommunities, we first evaluated the prevalence of preemptive competition [10, 20] among species. To do so, we evaluated community composition in summer of pools that only contained a single species in the previous spring. Our second objective was to test if the probability of observing a species pair in a pool is greater when the pool has more neighboring pools, and has experienced more frequent desiccation events.

To address our first objective, we retained only those pools that contained a single focal species in spring, and analyzed community composition in the subsequent summer. We identify four potential summer compositions: Same composition as in spring (1); Replacement of the focal species by one species (2); Replacement of the focal species by two species (3); Presence of two species, including the focal species (4). We modeled the probability for case (1) to occur, since if case (1) would be most frequent for each of the three focal species, this would be evidence for preemptive competition. We only used data of pools that were relatively isolated, i.e. had less than 20 surrounding pools within a 100 m radius, assuming these would better reflect the outcome of local dynamics. To these data, we fitted a Bayesian binomial generalized linear mixed model (GLMM) with a logit link function for each species. The response variable was binary: 0 (focal not present as only species in summer) or 1 (focal present as only species in summer). We modeled *p*_*A,ijt*_, the probability of observing the focal species *A* as the only species in the *j*^th^ pool of the *i*^th^ island in summer of year *t*. The fixed effect was the presence of the focal species as the single species in spring of the same year and in the same pool. We included island, pool identity (nested within island), and year as random effects:

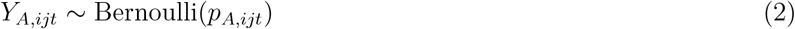

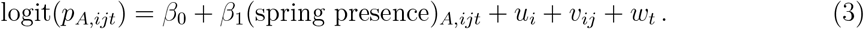

Where 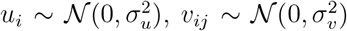, and 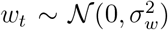. We relied on the default priors offered by the `brms` [43] package: improper uniform (flat) functions for all means, and half-Student t-distributions for all standard deviations. The latter always had 3 degrees of freedom, location parameter zero, and scale parameter equal to 2.5 times the standard deviation of the response variable after applying the logit link function.

To address our second objective, an intuitive approach would have been to use the number of pools within a cluster as a predictor of the fraction of pools where species coexist. However, this has as a caveat that pool density differed per cluster, possibly introducing important differences in dispersal rates between them. We therefore instead refocused our analysis on the pool level, asking if co-occurrence was more likely in pools surrounded by a greater number of other pools and in clusters with higher desiccation rates (and therefore presumably higher dispersal rates). Our statistical method consisted of fitting a Bayesian binomial generalized linear mixed model (GLMM) with a logit link function. The response variable was binary: 0 (single species present in a pool) or 1 (pair present in a pool). We modeled *p*_*ijkt*_, the probability of observing a species pair in the *j*^th^ pool on the *i*^th^ island, belonging to cluster *k*, in year *t*. Fixed effects were the number of surrounding pools (*P*_*ij*_; constant across years for each pool) and the simulated desiccation rate (*S*_*kt*_; varying among clusters and years). We included island, pool identity (nested within island), and year as random effects to account for the hierarchical structure of the sampling design:

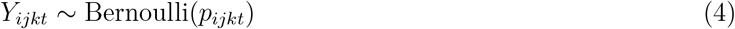

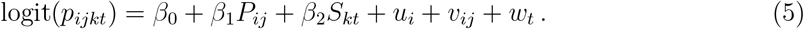

Where 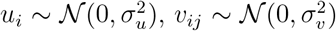, and 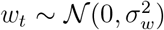. We fitted this model for both the spring and summer samples and for two distances around a pool (*b* = 50 m and 100 m). As for our first objective, we used the R [35] package `brms` [43] for model fitting, with the same default priors as before.

### 2.4 Computer code and data

All code was written in the free software R [35], version 4.4.0 Puppy Cup, using the packages `bayesplot, rootSolve, feasoverlap, geodist, gridExtra, brms, patchwork, reticulate, xtable,` and `tidyverse` [36, 37, 43–51], all available from CRAN.

## 3 Results

### 3.1 Species persistence

We start our analysis by partitioning the probability that a randomly chosen focal species *i* persists in a random patch *l* (i.e., that is has a density 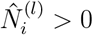) into two ecologically relevant cases, using the law of total probability:

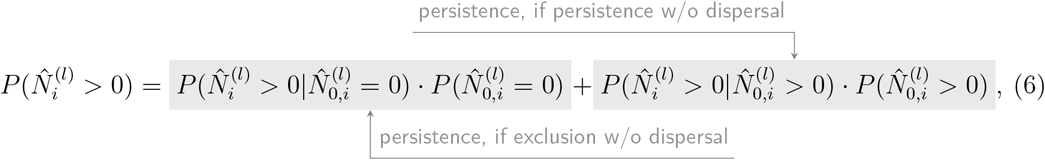

where the subscripts 0 denote densities that species *i* would attain if patches were completely disconnected and thus there was no dispersal. The first part of this probability deals with persistence in case the focal species would be excluded without dispersal; the second part deals with persistence when it would have persisted without dispersal.

Focusing on the first part, we are able to express species *i*’s equilibrium in a patch *l* solely as a function of its equilibrium without dispersal (Supporting Information, Section 1.2.1):

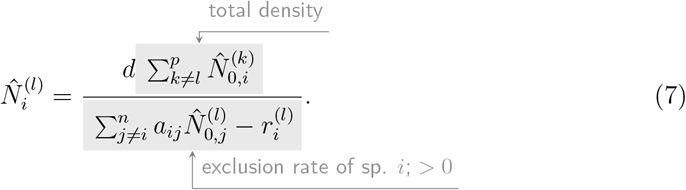

The numerator of Eq. 7 contains the sum of species *i*’s equilibrium densities (in absence of dispersal) across all patches except the focal patch. The denominator of Eq. 7 is the negative of the growth rate when species *i* invades patch *l* when all other species are at their equilibrium if there were no dispersal—that is, the invasion growth rate of coexistence theory [32]. The invasion growth rate quantifies the growth rate of a species following their introduction in an established community and must be positive for the species to persist. Given the case at hand (exclusion without dispersal), the invasion growth rate must be *negative*, which is why we coin its negative as the *exclusion rate* of species *i*. This means Eq. 7’s denominator is necessarily positive. Thus, we recover the result that a species is guaranteed to persist in all patches if dispersal is positive (*d* > 0) and when it would survive without dispersal in at least one other patch that is part of the spatial network. This conclusion is general: it does not depend on the exact parameter setting of Eq. 1 and therefore should hold for any number of species, and any species interaction scheme or landscape structure.

In realistic settings, having a positive density is insufficient for long-term persistence: low densities increase the risk of stochastic extinction. Thus, in many real-world applications, 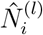 needs to be larger than some threshold. Eq. 7 shows that the probability for this to happen is larger when the focal species is more abundant across the spatial network of disconnected patches (has greater 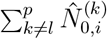), dispersal is larger (*d* is larger), the spatial network is larger (greater *p*), and the species gets excluded more slowly. This result quantifies how species persistence emerges as the net result of regional abundance and local dynamics, both assessed in the case patches are disconnected. Importantly, more challenging local interactions can be compensated by adding patches, i.e., increasing the size of the spatial network.

By making strong assumptions on the distribution of the growth rates (regional equivalence, see Methods) and species interactions (diffuse interactions that are weaker than intraspecific interactions, see Methods), we can calculate exactly how the numerator of Eq. 7 (total density across patches) depends on interaction strength and landscape size (Fig. 1A), and how the denominator of Eq. 7 (exclusion rate) depends on the interaction strength *a* (Fig. 1B) (Supporting Information, Section 1.2.2 and 1.2.3). Combining both parts of Eq. 7 shows how local persistence of a focal species depends on landscape size, dispersal rate, and local interaction strength (Fig. 1C). Persistence is more likely in larger and better-connected networks, and when interactions are weaker, regional density is higher, and exclusion is slower, respectively.

**Figure 1:**
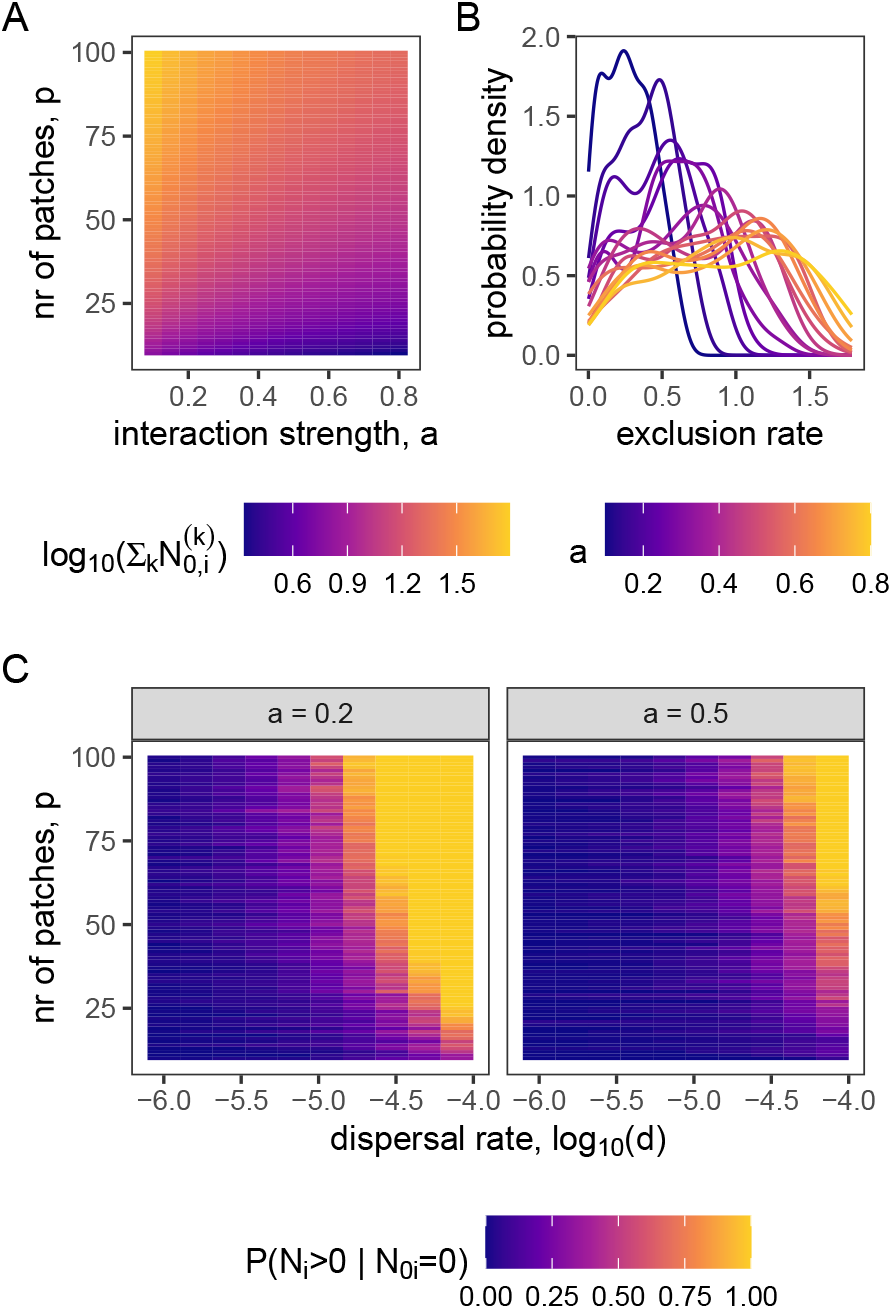
Landscape size and interaction strength predict species persistence 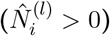 in case exclusion would happen without dispersal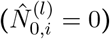. 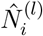 and 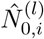 are equilibrium densities with and without dispersal, respectively. A: In case interspecific interactions are fixed to *a*, a species’ total density across all patches is also fixed, identical for all species, and only depends on *a* and the number of patches *p*. B: Species’ exclusion rates vary across patches and become larger with increasing *a*. C: Weak dispersal promotes the probability to persist, and more so when interactions are weak and the network is large.

We finally focus on the second case of Eq. 6, dealing with persistence of species that persist without dispersal. In the Supporting Information, Section 1.3, we present evidence that whenever a species persists without dispersal, it almost certainly persist with dispersal as well: 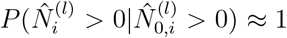 whenever species are regionally equivalent, and interspecific interactions are diffuse and weaker than intraspecific interactions.

### 3.2 Coexistence: Community-level patch occupancy

We can now use our species-level results to probe community-level patch occupancy. We start by expanding Eq. 6 to the community-level, and obtain the probability that all species persist in a randomly drawn patch:

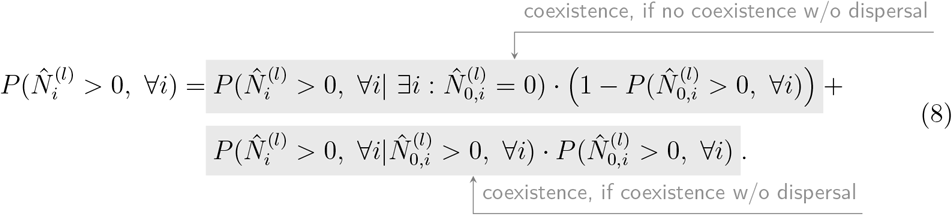

For the first case (no coexistence without dispersal), our species-level results imply that the *m* species that do persist without dispersal will persist with dispersal as well. We can therefore focus on the *n* − *m* species that do not persist without dispersal. We approximate the probability for all of these to persist as the (*n* − *m*)th power of 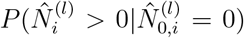, where 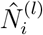 is given by Eq. 7. Denoting the distribution across the patches of the number of persisting species as *f* (*m*) (as done in Supporting Information, Section 1.2.2), we write:

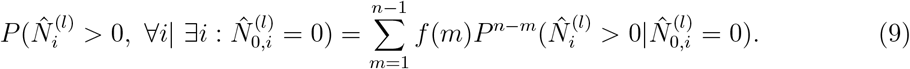

For the second case (coexistence without dispersal), our species-level results imply that 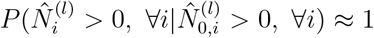 (Supporting Information, Section 1.3). Overall, we obtain the final result for patch occupancy:

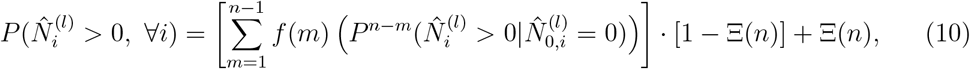

where we have abbreviated 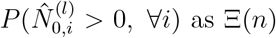, which is the feasibility domain size, a quantity used in coexistence theory to study the scope for coexistence when growth rates vary [29, 30]. For the case of regional equivalence and diffuse interactions that are weaker than intraspecific interactions, we can compute patch occupancy analytically. The calculated values compare well with dynamical simulations of Eq. 1 (Fig. 2). Specifically, patch occupancy sharply increases with the dispersal rate *d*, and this happens at lower *d* when interactions are weaker or the spatial network is larger.

**Figure 2:**
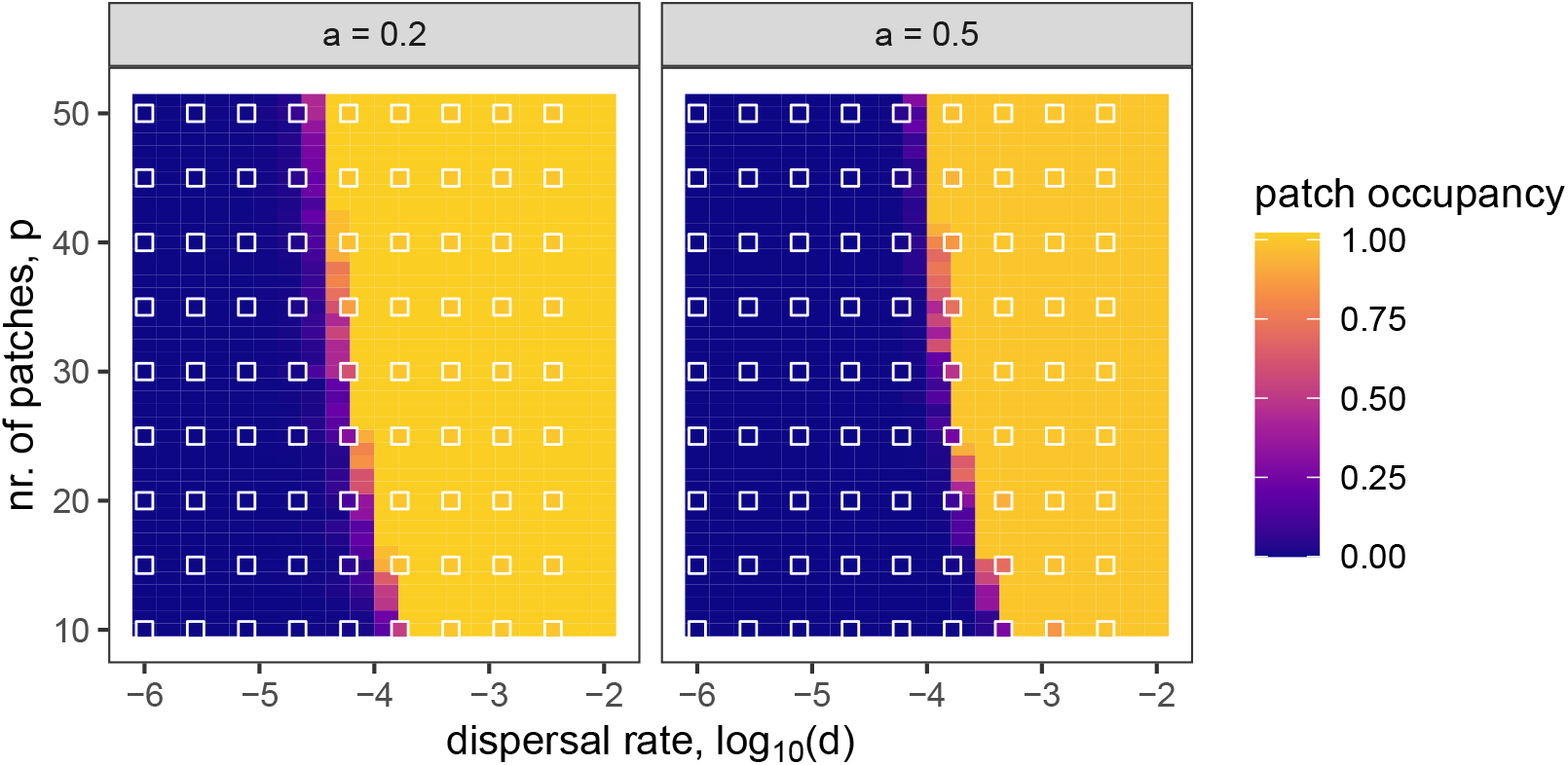
Predicted (background color) and simulated (filled squares, overplotting the heat map in the background) community-level patch occupancy (fraction of patches in which all regionally available species persist) as a function of dispersal rate and landscape size. Simulations are numerical integrations of Eq. 1 with *n* = 10 species. The detailed simulation protocol is in Section 2.2. Background colors are predictions for *n* = 10 by Eq. 10. Panels are two values of interspecific interaction strength *a*, which is the same across all species pairs (diffuse interactions) and always weaker than intraspecific interactions (all set to 1). Note that numerical exploration suggests values up until ≈ 0.005 are weak.

Our analytical predictions assume that only intrinsic growth varies spatially, that species are regionally equivalent, and that competition is weaker than intraspecific interactions and diffuse. Our analyses also imply that any pair of patches are equally well-connected, which has allowed us to use a single dispersal rate *d*. Finally, our results only use equilibrium population densities to determine coexistence, but the stability of these equilibria also matters for long-term coexistence. We therefore ran dynamical simulations where we seeded species across patches, allowed interactions to vary across patches and across species pairs within a patch (abandoning diffuse interactions), considered various mean interaction strengths (including preemptive competition), allowed a single species to have a higher regional mean intrinsic growth rate than the others (breaking regional equivalence of the growth rate), and allowed dispersal to occur according to explicit dispersal kernels. Adding all these complexities did not qualitatively change the results: weak dispersal always promotes coexistence, and more so in larger landscapes. Only when dispersal becomes sufficiently strong (corresponding to ≈ 0.01, Supporting Information, Fig. . S1), deviating from the premise of our analysis, and for strong competition (*a* = 0.5 or 1), the effect of dispersal on patch occupancy becomes unimodal (Fig. 3). Under these conditions, the effect of landscape size, if any, is consistently positive.

**Figure 3:**
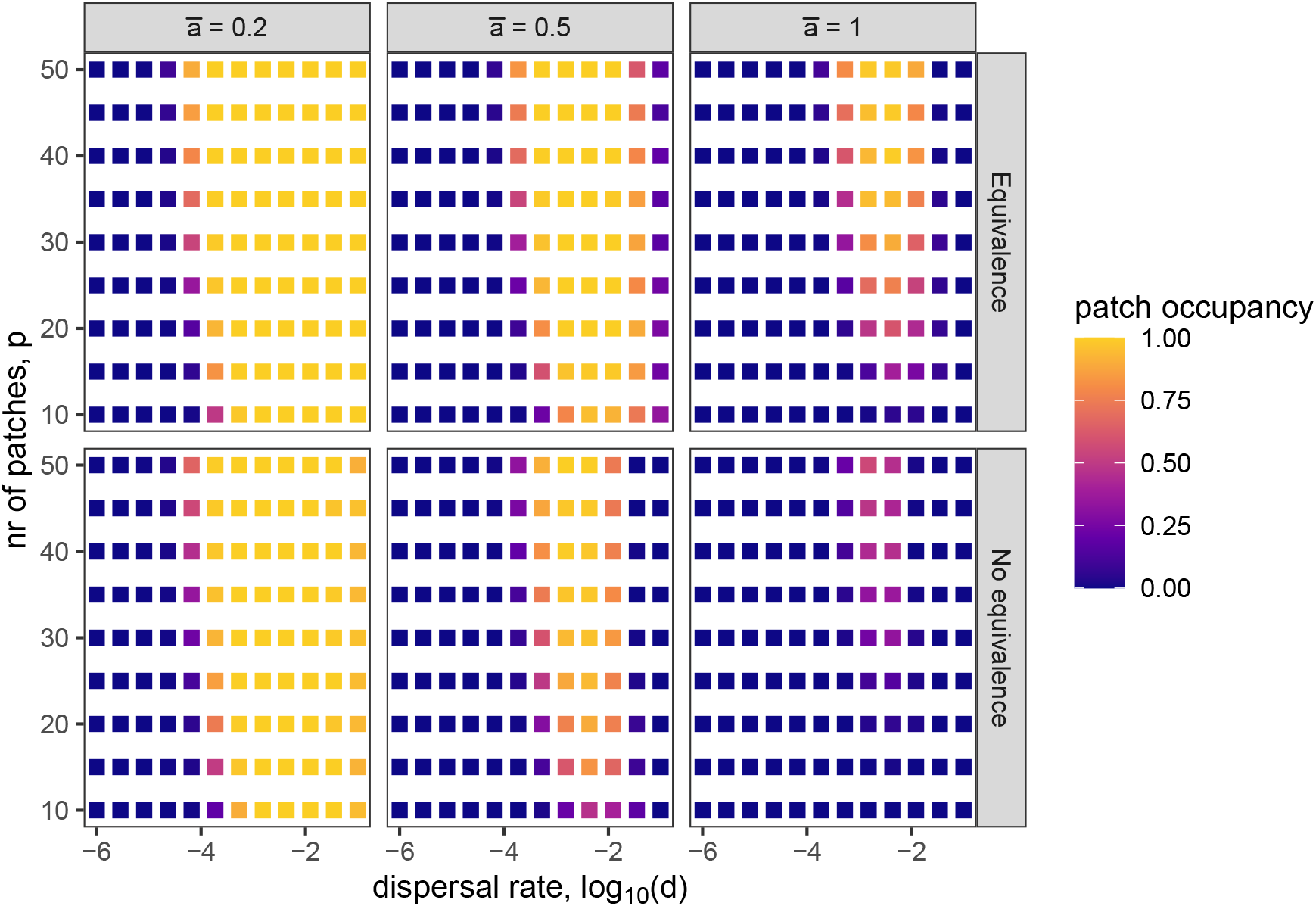
Weak dispersal and landscape size promote community-level patch occupancy, even when relaxing the assumptions of our analyses. Filled squares are simulations where species interactions vary spatially, and these interactions are no longer diffuse but vary across species pairs within a patch, with CV(*a*_*ij*_) = 0.2. Dispersal is no longer implicit but follows an exponential kernel with characteristic distances of species that are evenly spaced between 4 and 0.5 (distances between patches vary between about 10^−5^ and 1.4). Furthermore, there is either regional equivalence (all species have the same regional mean growth rate; upper row), or regional equivalence (species have different regional mean growth rate; bottom row). Note that patch occupancy declines when dispersal is too fast (> 0.01) and interactions are strong (*ā* = 0.5 or 1). As in the other figures, *n* = 10.

### 3.3 *Daphnia* co-occurrence in rockpools

For all species, we found that if it was present alone in a rockpool in spring, it was very likely present alone in that same rockpool in summer as well (probability of about 90%, Fig. 4). Specifically, the regression model showed that having the focal species as the only species in a pool in spring made it about 1000 times more likely to have that same species present as the only species in summer too. Supporting Information, Fig. S2 shows that predicted matched observed response variable distributions.

**Figure 4:**
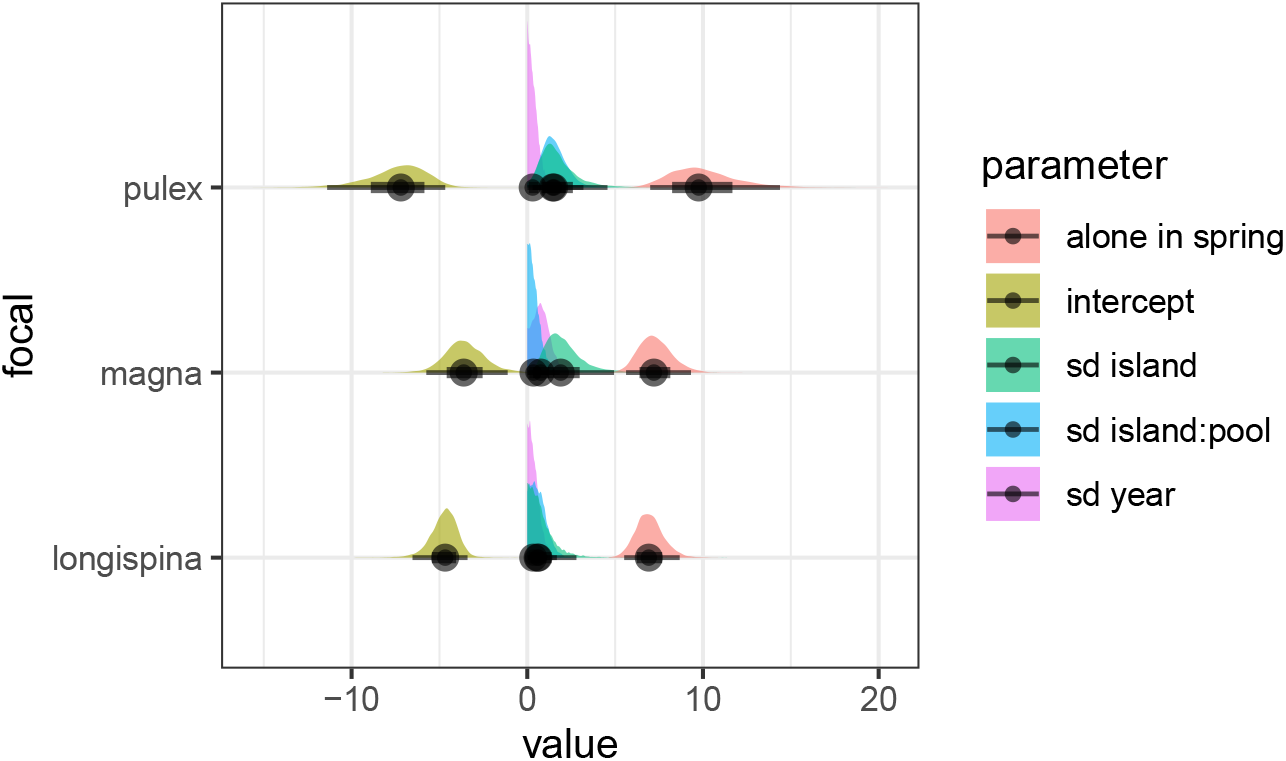
Evidence for preemptive competition across species. Posterior distributions of key model parameters from Bayesian hierarchical models fitted separately for each focal species. Positive estimates for the effect of being alone in spring for all three species indicate preemptive competition. Distributions show posterior uncertainty, with points denoting posterior means and thick intervals representing credible intervals; random-effect standard deviations quantify variation among islands, pools, and years. Values are in Supporting Information, Table S1.

Co-occurrence of two *Daphnid* species was rare in pools surrounded by less than 10 other pools. Only 20% of those pools had species pairs. However, this percentage increased with the number of surrounding pools, regardless of the considered distance around a pool (Fig. 5). This effect was consistent between sampling time (spring or summer). In contrast, the desiccation rate did not significantly influence co-occurrence. Supporting Information, Fig. S3 shows the posterior distributions, while Supporting Information, Fig. S4 confirm that the posterior predictive distribution matches the observed response variable distributions well.

**Figure 5:**
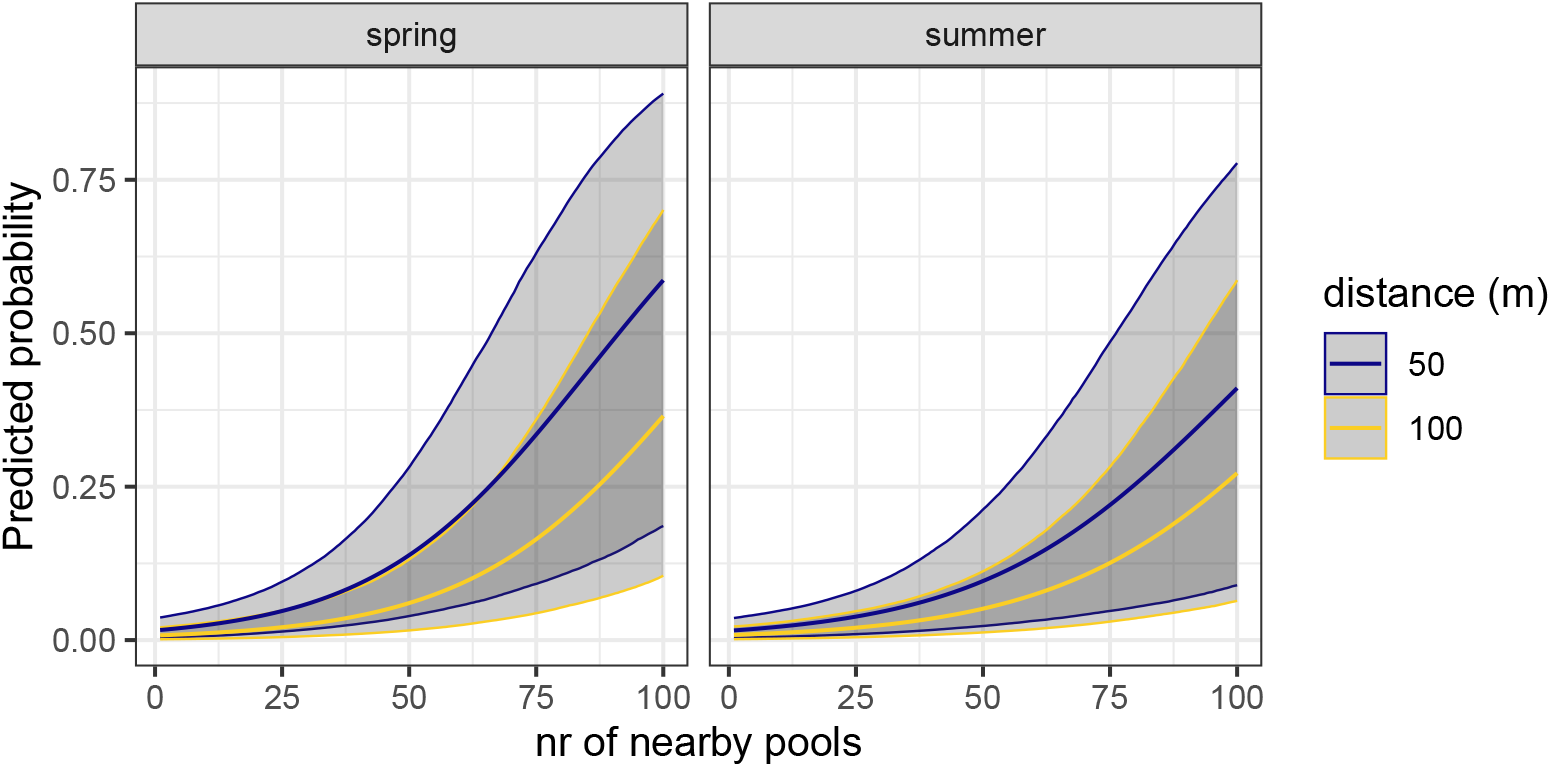
The number of surrounding pools increases occurrence probability of pairs. Predicted probability of occurrence as a function of the number of nearby pools, shown separately for spring and summer and for two spatial scales (50 m and 100 m). Lines represent posterior mean predictions from the Bayesian hierarchical model, with shaded ribbons indicating 95% credible intervals. Posterior distributions are plotted in Supporting Information, Fig. S3 and printed in Supporting Information, Table S2.

## 4 Discussion

We have presented theory and simulations showing that weak dispersal and landscape size (number of patches) inevitably promote local persistence and coexistence of competing species. Our analyses show how persistence emerges as a balance between local extinction and the regional density of a species, and that both parts can be estimated by treating the patches as if they were disconnected. We show that the fraction of patches in which all regionally available species coexist, which we coin *community-level patch occupancy*, is tightly linked to the size of the feasibility domain [29, 30], an important concept in coexistence theory. Finally, we present data of a microcrustacean metacommunity inhabiting rock pools at the coast of Finland, showing that species more often co-occur when they are surrounded by more patches (rock pools).

Our results shed new light on classic findings on biodiversity maintenance in spatial networks, obtained with a variety of models and data collection methods. The equilibrium theory of island biography [52] and its heir, the neutral theory of biodiversity [53, 54], both predict that local richness emerges as a balance between local extinction and immigration. Our results highlight how the negative of the invasion growth rate drives local extinction. We note that the *extinction rate* can be negative either because of unsuitable local conditions (environmental filtering, 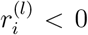, [55]), or because of too intense interspecific competition (large 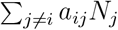). Our results suggest that immigration, when dispersal is weak, is a simple linear function of the size of the landscape: all else equal, total regional density is smaller in smaller networks, which makes density in a focal patch to fall more readily below the threshold for extinction. Explanations for the positive effect of landscape size on local richness often evoke a greater niche availability in larger networks [56]. Our results show that landscape size in and by itself can have an effect on species persistence, and therefore mean species richness, without greater niche availability. In spirit, this mechanism is the same as the one driving the species-area relationship in the equilibrium theory of island biogeography and its recent extensions, which states that smaller islands host fewer species because population densities are lower, making stochastic extinctions more likely [52, 57].

Our results can also be interpreted in the context of more recent concepts in spatial population and community ecology. At the level of single metapopulations, a classic result is that a species can only persist when its metapopulation capacity *λ* exceeds a well-defined threshold value [24–27]. The metapopulation capacity is obtained (at least in simple cases) as the leading eigenvalue of the matrix *M* ^(*kl*)^*/E*^(*k*)^, where *M* ^(*kl*)^ encodes the dispersal rate from patch *l* to patch *k*, and *E*^(*k*)^ is the rate of extinction in patch *k*. All other things equal, adding more patches to a landscape necessarily leads to a higher *λ* [25, 27]. Thus, Eq. 7 reflects the same idea as the fact that *λ* must exceed a threshold value, with the dispersal matrix *M* ^(*kl*)^ and extinction rates *E*^(*k*)^ playing the role of Eq. 7’s numerator and denominator, respectively. Our main result can therefore be interpreted as a community-level extension of the well-known criterion for metapopulation persistence. This goes beyond the earlier results of Häussler et al. [26] in that it applies to all communities described by Eq. 1, not just trophic metacommunities with strictly bottom-up interactions. More recently, Padmanabha et al. [58] have introduced the concept of local landscape-mediated fitness as the ratio between colonization and extinction at a focal patch, and show that persistence is more likely as this ratio increases. Again, this is conceptually similar to our results (Eq. 7).

At the level of multiple species, of particular interest are mass effects: this mechanism permits persistence of a species when dispersal overrules locally unfavorable conditions. It is typically found to operate in homogeneous environment when dispersal is strong [19]. In our analyses, we show that a species that does not persist in absence of dispersal (i.e., has a negative invasion growth rate) is more likely to achieve a viable density when dispersal *or* the number of patches is sufficiently high (Eq. 7). This matches the verbal definition of a mass effect [16, 59], except that dispersal need not be high.

A calibration of a patch occupancy model to the same *Daphnia* data has shown that temperature affects rates of colonization and extinction, and often unequally so among species [60], confirming earlier studies on the same system [42, 61]. Amid these changes in local species interactions and most likely deviations from regional equivalence, our analysis suggests that coexistence is more likely in pools that are surrounded by a greater number of neighbor pools, offering support for our theoretical result that coexistence in a patch is a number’s game despite changing the specific parameter settings of our model. Instead, we did not find effects of desiccation rate on co-occurrence. This can be because of two reasons. First, our analyses suggest preemptive competition (Fig. 4) and that coexistence is rare in isolated pools (80% of those pools carry a single species only). Both of these findings point to strong interspecific competition, in which case the effect of dispersal is unimodel and hence harder to pick up (cf. Fig. 3). A second reason could be that dispersal did not vary sufficiently to elicit effects on co-occurrence. Since we do not know the exact relationship between desiccation and dispersal (as dispersal was not directly measured), we cannot exclude this possibility.

Our prediction that weak dispersal and landscape size are bound to promote biodiversity is supported by studies on microbiome composition, finding that greater host density (i.e., a higher number of patches), or more intense social interactions promote alpha diversity of the microbiome. This has been found for density of rodent hosts [62], grooming relationships among baboon hosts [63], and relationships between spouses in humans [64].

While our analyses and simulations together cover a wide range of ecological settings, we identify several avenues for future research. First, our analyses only focus on feasibility (having positive population density) as a proxy for persistence and coexistence, disregarding potential effects of landscape architecture on the stability of the metacommunity [65–67]. Available studies have found that dispersal has a positive effect on metacommunity stability when it is weak, and that landscapes with a greater number of patches are more stable.

These results show that our findings on population densities are unlikely to be disrupted by stability considerations. However, we considered communities where species do not cluster into functional groups, trophic levels, or different dispersal syndromes. In metacommunities with strong segregation, it is known that dispersal can be destabilizing too [68]. While our mathematical analyses are general for the case where species go extinct without dispersal (Eq. 7), they are less so for the case where species persist without dispersal. That is because, in the latter case, we had to assume diffuse interactions and regional equivalence to prove species persistence with dispersal when species persist without dispersal.

Second, we have ignored the stochastic nature of immigration and of local dynamics and only mimic some of its consequences by considering species with a too low density as extinct. Stochastic dynamics are an important component of community dynamics in general and of metacommunity dynamics in particular [19], and models of recurrent stochastic immigration have been shown to recreate well-documented macroecological patterns in models much like the one we use here [33].

Third, we have focused on local persistence and coexistence. Future efforts could extend our analyses to the regional scale, following earlier approaches to summarize all local dynamics into one framework to predict whether species can regionally coexist, using regional proxies for species interactions, growth, and dispersal [33, 60, 69].

Fourth, we assume there is no cost to dispersal, because in our model intrinsic growth rates do not depend on dispersal rates. This assumption likely does not hold [70] and is expected to reduce persistence by decreasing regionally available density (Numerator of Eq. 7), since total density in a patch scales with the mean intrinsic growth rate in that patch. Our results show that such reductions of the mean intrinsic growth rate are able to offset the beneficial effects of a greater number of patches *p* or a higher dispersal rate *d*, confirming earlier modeling results [71].

Fifth, we have focused on heterogeneous landscapes by letting species grow differently in different patches. Analyses of models for homogeneous landscapes (the growth rate of a species is the same across all patches) can make dispersal disruptive for persistence [19, 58]. However, species growth rates need not be spatially variable to have large spatial variation in local densities, which is what needed to make Eq. 7’s numerator positive for most species and thus promote persistence. Pettersson and Jacobi have shown that large variation of interaction strengths among species pairs induces spatial heterogeneity by itself, even when these interactions and intrinsic growth rates do not vary spatially [22]. In this scenario, multistable population dynamics can by themselves create spatial heterogeneity, which increases local richness when combined with weak dispersal. This confirms previous assertions that multistable dynamics combined with weak dispersal should stimulate local biodiversity [3].

Overall, our results support the inevitability of local persistence and coexistence of competing species in spatial networks, provided dispersal is weak. These results provide a stepping stone towards more complex scenarios of landscape structure, dispersal syndromes, and alternative species interactions schemes.

## 5 Acknowledgements

We thank Axel Rossberg for useful feedback on an earlier version of our work. This research used resources of the “Plateforme Technologique de Calcul Intensif (PTCI)” (http://www.ptci.unamur.be) located at the University of Namur, Belgium, which is supported by the FNRS-FRFC, the Walloon Region, and the University of Namur (Conventions No. 2.5020.11, GEQ U.G006.15, 1610468, RW/GEQ2016 et U.G011.22). The PTCI is member of the “Consortium des Équipements de Calcul Intensif (CÉCI)” (http://www.ceci-hpc.be).

## 6 Author contributions

Conceptualization: FDL, AG, GB Methodology: FDL, GB Software: FDL Formal analysis: FDL, GB Data curation: OB, DE Writing – original draft: FDL Writing – review & editing: all authors

## 8 Open research statement

Data and computer code are available online at https://doi.org/10.5281/zenodo.15311497 and https://github.com/fdelaend/occupancy.

