## Supplemental Information for "Weak dispersal and landscape size inevitably promote local biodiversity in heterogeneous metacommunities of competing species"

De Laender et al.

### 1 Supplemental methods

#### 1.1 Weak dispersal approximation

Let us write the equilibrium condition of the main model in the main text by setting the time derivative on its left hand side to zero:

$$\hat{N}_i^{(l)} \left( r_i^{(l)} - \hat{N}_i^{(l)} - \sum_{j \neq i}^{n} a_{ij} \hat{N}_j^{(l)} - (p-1)D \right) + D \sum_{k \neq l}^{p} \hat{N}_i^{k} = 0. \tag{1}$$

We make one further change: the terms containing a factor of $D$ are multiplied and divided by a constant $D_0$ expressing unit dispersal. This leaves the expression unchanged, but allows one to write solutions in terms of $D/D_0$, which is a unitless parameter. Let us denote this fraction by $d$; then $D = D_0 d$. Eq. 1 can now be written as

$$\hat{N}_i^{(l)} \left( r_i^{(l)} - \hat{N}_i^{(l)} - \sum_{j \neq i}^{n} a_{ij} \hat{N}_j^{(l)} - (p-1)D_0 d \right) + D_0 d \sum_{k \neq l}^{p} \hat{N}_i^{k} = 0. \tag{2}$$

We assume that $0 \leq d \ll 1$. In that case, the solution to Eq. 2 can be written as a power series in $d$ (which is why it was important to make it dimensionless):

$$\hat{N}_i^{(l)} = \hat{N}_{0,i}^{(l)} + d\hat{N}_{1,i}^{(l)} + d^2 \hat{N}_{2,i}^{(l)} + \ldots \tag{3}$$

We only keep the constant and linear terms in $d$, ending up with the approximation

$$\hat{N}_i^{(l)} \approx \hat{N}_{0,i}^{(l)} + d\hat{N}_{1,i}^{(l)}. \tag{4}$$

We substitute Eq. 4 into Eq. 2 up to that linear term:

$$\begin{aligned} \left( \hat{N}_{0,i}^{(l)} + d\hat{N}_{1,i}^{(l)} \right) \left( r_i^{(l)} - \left( \hat{N}_{0,i}^{(l)} + d\hat{N}_{1,i}^{(l)} \right) - \sum_{j \neq i}^{n} a_{ij} \left( \hat{N}_{0,j}^{(l)} + d\hat{N}_{1,j}^{(l)} \right) - (p-1)D_0 d \right) \\ + D_0 d \sum_{k \neq l}^{p} \left( \hat{N}_{0,i}^{(k)} + d\hat{N}_{1,i}^{(k)} \right) = 0. \end{aligned} \tag{5}$$

Since $d$ is a parameter and the equation must hold for any value it might take, this equation can only hold if the 0th-order terms and the 1st-order terms in $d$ both individually sum to zero. Collecting the 0th-order terms first:

$$\hat{N}_{0,i}^{(l)} \left( r_i^{(l)} - \hat{N}_{0,i}^{(l)} - \sum_{j \neq i}^{n} a_{ij} \hat{N}_{0,j}^{(l)} \right) = 0, \tag{6}$$

which is the usual Lotka–Volterra equilibrium condition in patch $l$.

Continuing with the 1st-order terms in $d$ in Eq. 5:

$$\hat{N}_{0,i}^{(l)} \left( -d\hat{N}_{1,i}^{(l)} - \sum_{j \neq i}^{n} d a_{ij} \hat{N}_{1,j}^{(l)} - (p-1) D_0 d \right) + d\hat{N}_{1,i}^{(l)} \left( r_i^{(l)} - \hat{N}_{0,i}^{(l)} - \sum_{j \neq i}^{n} a_{ij} \hat{N}_{0,j}^{(l)} \right) + D_0 d \sum_{k \neq l}^{p} \hat{N}_{0,i}^{(k)} = 0. \tag{7}$$

Dividing by $d$:

$$\hat{N}_{0,i}^{(l)} \left( -\hat{N}_{1,i}^{(l)} - \sum_{j \neq i}^{n} a_{ij} \hat{N}_{1,j}^{(l)} - (p-1) D_0 \right) + \hat{N}_{1,i}^{(l)} \left( r_i^{(l)} - \hat{N}_{0,i}^{(l)} - \sum_{j \neq i}^{n} a_{ij} \hat{N}_{0,j}^{(l)} \right) + D_0 \sum_{k \neq l}^{p} \hat{N}_{0,i}^{(k)} = 0. \tag{8}$$

Rearranging and simplifying:

$$\hat{N}_{1,i}^{(l)} \left( r_i^{(l)} - 2\hat{N}_{0,i}^{(l)} - \sum_{j \neq i}^{n} a_{ij} \hat{N}_{0,j}^{(l)} \right) - \hat{N}_{0,i}^{(l)} \sum_{j \neq i}^{n} a_{ij} \hat{N}_{1,j}^{(l)} + D_0 \left( \sum_{k \neq l}^{p} \hat{N}_{0,i}^{(k)} - (p-1) \hat{N}_{0,i}^{(l)} \right) = 0. \tag{9}$$

This is a linear equation in the $\hat{N}_{1,i}^{(l)}$, assuming that the $\hat{N}_{0,j}^{(l)}$ have already been calculated and are known. In that case, the bracketed term above is the coefficient of $\hat{N}_{1,i}^{(l)}$ and $-a_{ij}\hat{N}_{0,i}^{(l)}$ are the coefficients of $\hat{N}_{1,j}^{(l)}$, $\forall\ j \neq i$, and the final term $D_0(\sum_{k \neq l}^{p} \hat{N}_{0,i}^{(k)} - (p-1)\hat{N}_{0,i}^{(l)})$ is the intercept. In what follows, we use Eq. 9 to analyze persistence of a focal species $i$ in patch $l$ for two cases: either species $i$ goes extinct when there is no dispersal ($\hat{N}_{0,i}^{(l)} = 0$), or it persists when there is no dispersal ($\hat{N}_{0,i}^{(l)} > 0$).

Note that we omit $D_0$ in the next sections and in the main text, since $D_0 = 1$ is a unit dispersal rate introduced just for being able to express a power series in the unitless parameter $d = D/D_0$.

#### 1.2 Exclusion without dispersal

##### 1.2.1 Equilibrium density

If the focal species $i$ goes extinct in patch $l$ when there is no dispersal, it implies that $\hat{N}_{0,i}^{(l)} = 0$. Eq. 9 then simplifies to:

$$\hat{N}_{1,i}^{(l)} \left( r_i^{(l)} - \sum_{j \neq i}^{n} a_{ij} \hat{N}_{0,j}^{(l)} \right) + \sum_{k \neq l}^{p} \hat{N}_{0,i}^{(k)} = 0, \tag{10}$$

leading to:

$$\hat{N}_{1,i}^{(l)} = \frac{\sum_{k \neq l}^{p} \hat{N}_{0,i}^{(k)}}{\sum_{j \neq i}^{n} a_{ij} \hat{N}_{0,j}^{(l)} - r_i^{(l)}}, \tag{11}$$

which implies that:

$$\begin{aligned} \hat{N}_i^{(l)} &= \hat{N}_{0,i}^{(l)} + d\hat{N}_{1,i}^{(l)} \\ &= 0 + \frac{d \sum_{k \neq l}^{p} \hat{N}_{0,i}^{(k)}}{\sum_{j \neq i}^{n} a_{ij} \hat{N}_{0,j}^{(l)} - r_i^{(l)}} \\ &= \frac{d \sum_{k \neq l}^{p} \hat{N}_{0,i}^{(k)}}{\sum_{j \neq i}^{n} a_{ij} \hat{N}_{0,j}^{(l)} - r_i^{(l)}}. \end{aligned} \tag{12}$$

##### 1.2.2 Approximation for regional equivalence and diffuse interactions

Eq. 12 depends on two sums: the total density of the focal species $i$ across all disconnected patches but the focal patch $l$ ($\sum_{k \neq l}^{p} \hat{N}_{0,i}^{(k)}$), and the total density across all species within our focal patch $l$, in absence of dispersal ($\sum_{j \neq i}^{n} \hat{N}_{0,j}^{(l)}$). Because of regional equivalence, the first of these sums is the same for all species. It therefore suffices to know the total density across all species and all patches but the focal one, which we define here as $\mathcal{N}_0$, and then divide this number by the number of species:

$$\sum_{k \neq l}^{p} \hat{N}_{0,i}^{(k)} = \frac{\mathcal{N}_0}{n} \tag{13}$$

To compute $\mathcal{N}_0$, we need to know (1) how many patches contain how many species, and (2) what the total density is of a patch when we know its number of species.

To address (1), we first define $f(m)$ as the fraction of patches that have exactly $m$ locally coexisting species:

$$f(m) = \sum_s \xi(s) = \binom{n}{m} \Xi(m), \tag{14}$$

where $m \leq n$ and $\xi(s)$ is the feasibility domain size for a specific sub-community $\mathcal{C}_s$ of $m$ species ($|\mathcal{C}_s| = m\ \forall s$). The fact that all species pairs interact with the same strength $a$ implies that $\xi(s)$ is the same for all $\mathcal{C}_s$ and only depends on $m$: the feasibility domain shrinks with $m$, all else equal [1]. We define $\Xi(m)$ as the function describing this dependence.

To address (2), the total density across all $m$ species, where $m$ can take any value between 1 and $n$, we first rewrite Eq. 6 as a matrix equation: $\hat{\mathbf{N}}_0 = \mathbf{A}^{-1}\mathbf{R}$, where $\hat{\mathbf{N}}_0$ collects all equilibria, $\mathbf{A}^{-1}$ is the inverse of the interaction matrix, and $\mathbf{R}$ is the vector of growth rates. Given the diffuse interactions assumption, this interaction matrix is

$$\mathbf{A} = \begin{pmatrix} 1 & a & \dots & a \\ a & 1 & \dots & a \\ \dots & \dots & \dots & \dots \\ a & a & \dots & 1 \end{pmatrix}. \tag{15}$$

Because of the simple structure of $\mathbf{A}$, the Sherman-Morrison formula yields a particularly simple expression for $\mathbf{A}^{-1}$:

$$\mathbf{A}^{-1} = \mathbf{B}^{-1} - \frac{\mathbf{B}^{-1}\mathbf{u}\mathbf{v}^{\mathbf{T}}\mathbf{B}^{-1}}{1 + \mathbf{v}^{\mathbf{T}}\mathbf{B}^{-1}\mathbf{u}}, \tag{16}$$

where

$$\mathbf{B} = \begin{pmatrix} 1-a & 0 & \dots & 0 \\ 0 & 1-a & \dots & 0 \\ \dots & \dots & \dots & \dots \\ 0 & 0 & \dots & 1-a \end{pmatrix},$$
$$\mathbf{u} = \begin{pmatrix} \sqrt{a} & \sqrt{a} & \dots & \sqrt{a} \end{pmatrix}^{\mathbf{T}},$$
$$\mathbf{v} = \mathbf{u}^{\mathbf{T}}, \tag{17}$$

such that

$$\mathbf{B}^{-1} = \begin{pmatrix} 1/(1-a) & 0 & \dots & 0 \\ 0 & 1/(1-a) & \dots & 0 \\ \dots & \dots & \dots & \dots \\ 0 & 0 & \dots & 1/(1-a) \end{pmatrix},$$
$$\mathbf{B}^{-1}\mathbf{u}\mathbf{v}^{\mathbf{T}}\mathbf{B}^{-1} = \begin{pmatrix} a/(1-a)^2 & a/(1-a)^2 & \dots & a/(1-a)^2 \\ a/(1-a)^2 & a/(1-a)^2 & \dots & a/(1-a)^2 \\ \dots & \dots & \dots & \dots \\ a/(1-a)^2 & a/(1-a)^2 & \dots & a/(1-a)^2 \end{pmatrix},$$
$$1 + \mathbf{v}^{\mathbf{T}}\mathbf{B}^{-1}\mathbf{u} = 1 + ma/(1-a). \tag{18}$$

Therefore, the diagonal elements of $\mathbf{A}^{-1}$ are all identical and read

$$(a_{ij})^{-1}_{ii} = -\frac{a(m-2)+1}{(a-1)(a(m-1)+1)} \ \forall i, \tag{19}$$

while the off-diagonals, again all identical, read

$$(a_{ij})^{-1}_{ij} = \frac{a}{(a-1)(a(m-1)+1)} \ \forall j \neq i. \tag{20}$$

This means that $\hat{N}_{0,i}^{(l)}$ can now be computed as

$$\begin{aligned}\hat{N}_{0,i}^{(l)} &= (a_{ij})_{ii}^{-1} r_i^{(l)} + \sum_{j \neq i}^{m} (a_{ij})_{ij}^{-1} r_j \\ &= -\frac{a(m-2)+1}{(a-1)(a(m-1)+1)} r_i^{(l)} + \frac{a}{(a-1)(a(m-1)+1)} (m-1)\overline{r}_{m-1} \\ &= \frac{a(m-1)\overline{r}_{m-1} - r_i^{(l)}(a(m-2)+1)}{(a-1)(a(m-1)+1)},\end{aligned} \tag{21}$$

making the total biomass across all species, $\sum_j \hat{N}_{0,j}^{(l)}$:

$$\begin{aligned}\sum_j \hat{N}_{0,j}^{(l)} &= \sum_j \frac{a(m-1)\overline{r}_{m-1} - r_j^{(l)}(a(m-2)+1)}{(a-1)(a(m-1)+1)} \\ &= \frac{m\overline{r}_m}{1+a(m-1)},\end{aligned} \tag{22}$$

where $\overline{r}_{m-1}$ and $\overline{r}_m$ are the mean growth rates in a patch with $m-1$ and $m$ species, respectively. Because not all species persist in every patch (often, $m < n$), there will be a selection on $r_i^{(l)}$ [2]. More precisely, patches with $m < n$ tend to contain the $m$ species with the highest growth rates.

To approximate $\overline{r}_m$, we first define the probability distribution $g^{(l)}$ of the $r_i^{(l)}$ across all species within a patch $l$, and denote its mean as $\overline{r}$. If there is no dispersal, then for every species $i \notin \mathcal{C}$ (where $\mathcal{C}$ is again the composition in patch $l$, with $|\mathcal{C}| = m$, meaning $\mathcal{C}$ consists of $m$ elements; i.e., $m$ species persist), it must hold that

$$r_i^{(l)} < a \sum_j \hat{N}_{0,j}^{(l)}, \tag{23}$$

meaning $i$ can't invade into patch $l$. This implies that the mean growth rate of the species not persisting without dispersal in patch $l$ will be the mean of $g^{(l)}$, after truncation at $a \sum_j \hat{N}_{0,j}^{(l)}$. The distribution $g^{(l)}$ having a lower bound of zero, the mean of the excluded species, which we refer to here as $\overline{r}_{-m}$, will be located between zero and $a \sum_j \hat{N}_{0,j}^{(l)}$, i.e., $xa \sum_j \hat{N}_{0,j}^{(l)}$, with

$0 \leq x \leq 1$. We then obtain:

$$\begin{aligned} \overline{r}_{-m} &\approx xa \sum_j N_{0,j}^{(l)} \\ &\approx \frac{xam\overline{r}_m^{(l)}}{1+a(m-1)}, \end{aligned} \tag{24}$$

where we use Eq. 22 to obtain $a \sum_j \hat{N}_{0,j}^{(l)}$ and define the mean growth rate of the $m$ persisting species as $\overline{r}_m^{(l)}$. We can then substitute $\overline{r}_{-m}$ with $\frac{n\overline{r}-m\overline{r}_m^{(l)}}{n-m}$ and solve for $\overline{r}_m^{(l)}$, which leads to

$$\overline{r}_m \approx \frac{n\overline{r}(a(-m)+a-1)}{m(am(x-1)-anx+a-1)}. \tag{25}$$

Since $\overline{r}$ is constant across patches (i.e., same for all $l$) and known by design, we can approximate $\overline{r}_m$ (and therefore also $\overline{r}_{-m}$) using only the model parameters $a$, $m$, and $n$, and the parameter $x$. The accuracy of this approximation will depend on how well the selected $x$ allows estimating the mean of the truncated $g^{(l)}$. Setting $x = 0.5$ corresponds to a uniform distribution; here, we set $x = 0.45$ to account for the skew of $g^{(l)}$, which leads to predictions of $\overline{r}_m$ vs. $m$ that match simulated values (values taken from the simulation protocol outlined in the methods, subsetting to $d = 1e^{-6}$ to switch off dispersal):

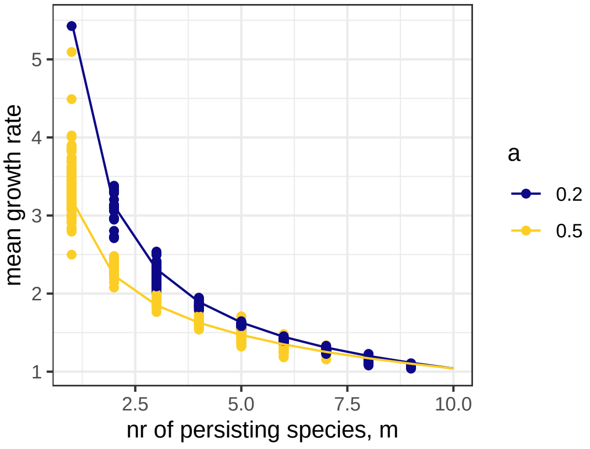

We then obtain $\mathcal{N}_0$ as the sum of the total densities across the $p-1$ patches with $m$ species (called $\mathcal{N}_{0,m}$), for $m$ varying from 1 to $n$:

$$\begin{aligned}\mathcal{N}_0 &= \sum_{m=1}^{n} \mathcal{N}_{0,m} \\ &= \sum_{m=1}^{n} f(m)\,(p-1)\,\frac{m\bar{r}_m}{1+a(m-1)},\end{aligned} \tag{26}$$

where Eq. 25 gives $\bar{r}_m$. Dividing by $n$ gives the total density across patches (per Eq. 13):

$$\sum_{k\neq l}^{p} \hat{N}_{0,i}^{(k)} = \frac{1}{n}\sum_{m=1}^{n} f(m)\,(p-1)\,\frac{m\bar{r}_m}{1+a(m-1)}. \tag{27}$$

In conclusion, Eq. 12's numerator ($d$ times total density across patches) is constant across patches and species, and only depends on interaction strength and network size. In contrast, Eq. 12's denominator (the exclusion rate, without dispersal) varies across patches and species. In case of diffuse interactions, it simplifies to $a\sum_{j\neq i}^{n} \hat{N}_{0,j}^{(l)} - r_i^{(l)}$, where $\sum_{j\neq i}^{n} \hat{N}_{0,j}^{(l)}$ is simply the total density across species in patch $l$. This sum follows a discrete probability distribution, where having a total density of a community with $m$ species (given by Eq. 22) happens with a probability that $m$ species are present, which is $f(m)$ (Eq. 14).

##### 1.2.3 Calculation of persistence probability

To calculate $P(N_i^{(l)} > 0 | \hat{N}_{0,i}^{(l)} = 0)$ we use Eq. 12. To calculate its numerator, $d\sum_{k\neq l} \hat{N}_{0,i}^{(k)}$, we use Eq. 27 and thus only need the model's basic parameters. To calculate its denominator, we sample 1000 values for $m$ with probability $f(m)$. Computing $f(m)$ involves computing the feasibility domain $\Xi(m)$ first (Eq. 14). To do so, one starts by taking the matrix of species interactions $\mathbf{A}$. For $m = n$, one can apply standard methods to get $\Xi(m)$. However, when $m < n$, we need to replace the columns of $\mathbf{A}$ that represent the absent species by unit vectors that have a $-1$ where the row number equals the column number [3]. Next, one can use this modified matrix to compute $\Xi(m)$ using the R package `feasoverlap` [4], constraining the feasibility domain on positive intrinsic growth rates.

Given the sampled $m$, we then calculate the 1000 corresponding values for the mean growth rate of persisting species ($\overline{r}_m^{(l)}$, using Eq. 25) and total local density ($\sum_{j\neq i}^{n} N_{0,j}^{(l)}$, using Eq. 22). We then sample $r_i^{(l)}$ such that $r_i^{(l)} < a\sum_{j\neq i}^{n} N_{0,j}^{(l)}$ from the distribution of growth rates $g^{(l)}$ (truncated normal, given in the main text). We then evaluate $P(N_i^{(l)} > 0|\hat{N}_{0,i}^{(l)} = 0)$ as the fraction of densities above the chosen threshold ($1e^{-3}$).

##### 1.3 Persistence without dispersal

For focal species persisting without dispersal, the per capita growth rate at equilibrium is zero. Assuming regional equivalence and diffuse interactions, we can write Eq. 9 as

$$
\begin{aligned}
0 &= \hat{N}_{1,i}^{(l)}\left(-\hat{N}_{0,i}^{(l)}\right) - a\hat{N}_{0,i}^{(l)}\sum_{j\neq i}^{n}\hat{N}_{1,j}^{(l)} + \sum_{k\neq l}^{p}\hat{N}_{0,i}^{(k)} - (p-1)\hat{N}_{0,i}^{(l)} \\
&= -\hat{N}_{1,i}^{(l)} - a\sum_{j\neq i}^{n}\hat{N}_{1,j}^{(l)} + \frac{\sum_{k\neq l}^{p}\hat{N}_{0,i}^{(k)}}{\hat{N}_{0,i}^{(l)}} - (p-1) \\
&= -\hat{N}_{1,i}^{(l)} - a\sum_{j\in\mathcal{C},j\neq i}\hat{N}_{1,j}^{(l)} - a\sum_{j\notin\mathcal{C}}\hat{N}_{1,j}^{(l)} + \rho_i - (p-1) \\
&= -\hat{N}_{1,i}^{(l)} - a\sum_{j\in\mathcal{C},j\neq i}\hat{N}_{1,j}^{(l)} - a\,(n-m)\,\overline{N}_1^{(l)} + \rho_i - (p-1),
\end{aligned}
\tag{28}
$$

where we obtained the second line by dividing all terms by $\hat{N}_{0,i}^{(l)}$, and define a variable $\rho_i = \sum_{k\neq l}^{p}\hat{N}_{0,i}^{(k)}/\hat{N}_{0,i}^{(l)}$ on the third line that quantifies the ratio between the total density of a species across all but the focal patch, and the local density of that species in the focal patch. Note that we expect that $E\left[\rho_i\right] \geq 0$. On the third line, we also split the sum $\sum_{j\neq i}^{n}\hat{N}_{1,j}^{(l)}$ in two parts: one part contains those species that persist without dispersal; the other part contains those that don't. Again, we introduce $\mathcal{C}$ as the community composition without dispersal in patch $l$ (with $|\mathcal{C}| = m$, meaning $m$ species persist without dispersal). On the fourth line, we define the mean $\overline{N}_1^{(l)}$ as the mean of all $\hat{N}_{1,j}^{(l)}$ taken across species that *do not* persist without dispersal in patch $l$. For reasons explained in Section 1.2.1, this mean cannot be negative.

Eq. 28 is a system of $m$ linear equations in the $\hat{N}_{1,i}^{(l)}$. The matrix holding the coefficients of this linear system has $-1$ on its diagonal and $-a$ as off-diagonals. The intercepts are in a vector with elements $-a\,(n-m)\,\overline{N}_1^{(l)} + \rho_i - (p-1)$. Because of the simple coefficient matrix, the inverse of the coefficient matrix has $\frac{a(m-2)+1}{(a-1)(a(m-1)+1)}$ on its diagonal, and $-\frac{a}{(a-1)(a(m-1)+1)}$ on all off-diagonals. The solution to this system is therefore

$$
\begin{aligned}
\hat{N}_{1,i}^{(l)} =& \left(\frac{a(m-2)+1}{(a-1)(a(m-1)+1)}\right)\left(-a\,(n-m)\,\overline{N}_1^{(l)} + \rho_i - (p-1)\right) \\
&+ \sum_{j\in\mathcal{C}, j\neq i}\left(-\frac{a}{(a-1)(a(m-1)+1)}\right)\left(-a\,(n-m)\,\overline{N}_1^{(l)} + \rho_j - (p-1)\right) \\
=& \left(\frac{a(m-2)+1}{(a-1)(a(m-1)+1)}\right)\left(-a\,(n-m)\,\overline{N}_1^{(l)} + \rho_i - (p-1)\right) \\
&+ (m-1)\left(-\frac{a}{(a-1)(a(m-1)+1)}\right)\left(-a\,(n-m)\,\overline{N}_1^{(l)} + \overline{\rho} - (p-1)\right) \\
=& \frac{a\left((a-1)\overline{N}_1^{(l)}(n-m) - m\overline{\rho} + m\rho_i + \overline{\rho} + p - 2\rho_i - 1\right) - p + \rho_i + 1}{(a-1)(a(m-1)+1)} \\
=& \frac{a\left((1-a)\overline{N}_1^{(l)}(n-m) + (m-1)\overline{\rho} + (2-m)\rho_i\right) + (p-1)(1-a) - \rho_i}{(1-a)(a(m-1)+1)},
\end{aligned}
\tag{29}
$$

where $\overline{\rho}$ must be nonnegative because the individual $\rho_i$ must be as well.

This means that the equilibrium density with dispersal is:

$$
N_i^{(l)} = \underbrace{N_{0,i}^{(l)}}_{>0\text{, because persistence w/o dispersal}} + d\,\frac{a\left(\overbrace{(1-a)\overline{N}_1^{(l)}(n-m)}^{>0} + \overbrace{(m-1)\overline{\rho}}^{\geq 0} + (2-m)\rho_i\right) + \overbrace{(p-1)(1-a)}^{>0} - \overbrace{\rho_i}^{\geq 0}}{\underbrace{(1-a)(a(m-1)+1)}_{>0}},
\tag{30}
$$

where annotations highlight terms in Eq. 30 that will always be positive, if interspecific interactions are weaker than intraspecific interactions ($a < 1$). The sign of $N_i^{(l)}$ will depend on the value of $m$ and $\rho_i$, however. It is helpful to consider two limiting cases: one where interactions are very strong ($a \to 1$), and one where they are very weak ($a \to 0$). When interactions are very strong, every patch only contains a single species, i.e. the one with the highest growth rate $r_i^{(l)}$. If we focus on the patch where the focal species is present, this implies that it must be absent from the other patches, making $\rho_i = 0$. This cancels out all terms with $\rho_i$ in Eq. 30, making $N_i^{(l)} > 0$. When interactions are very weak, all species occur nearly everywhere, making $\rho_i \approx p-1$ and $m \approx n$, making Eq. 30:

$$\begin{aligned} N_i^{(l)} &\approx N_{0,i}^{(l)} + d\frac{a\left((n-1)(p-1)+(2-n)(p-1)\right)+(p-1)(1-a)-(p-1)}{(1-a)(a(n-1)+1)} \\ &= N_{0,i}^{(l)} + d\frac{a\left(p-1\right)+(p-1)(1-a)-(p-1)}{(1-a)(a(n-1)+1)} \\ &= N_{0,i}^{(l)}, \end{aligned} \tag{31}$$

which confirms that $P(N_i^{(l)} > 0 | N_{0,i}^{(l)} > 0) \approx 1$: if a species persists without dispersal, it will almost certainly persist with dispersal as well.

155 We numerically calculated Eq. 29 using similar methods as in Section 1.2.3: sampling $m$ from $f(m)$, calculating all the terms from 29, and finally calculating $N_i^{(l)}$ for all sampled values, and evaluating which fraction is greater than the chosen extinction threshold of $1e^{-3}$. These numerical calculations confirm that species persisting without dispersal will almost certainly persist with dispersal as well:

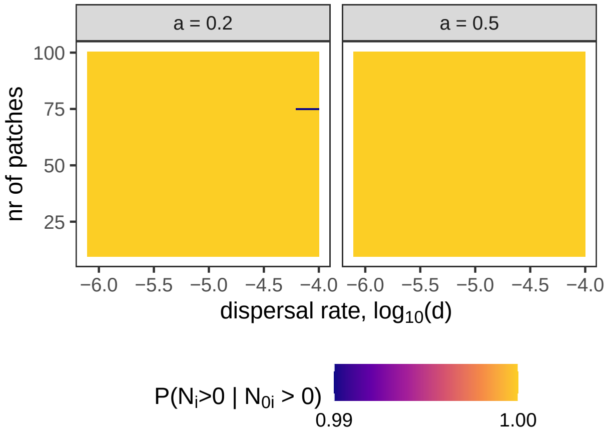

### 2 Supplemental results: statistical models of the *Daphnia* data

#### 2.1 Preemptive competition

Table S1: Estimated coefficients of the models (one per focal species), where *group* refers to the random effects.

| effect | component | group | term | estimate | std.error | conf.low | conf.high | focal |
|---|---|---|---|---|---|---|---|---|
| fixed | cond | | (Intercept) | -3.57 | 1.15 | -5.75 | -1.12 | magna |
| fixed | cond | | as.factormagna_spring1 | 7.28 | 0.94 | 5.62 | 9.32 | magna |
| ran_pars | cond | island | sd__(Intercept) | 2.11 | 1.10 | 0.63 | 4.94 | magna |
| ran_pars | cond | island:pool | sd__(Intercept) | 0.43 | 0.34 | 0.02 | 1.26 | magna |
| ran_pars | cond | year | sd__(Intercept) | 0.80 | 0.48 | 0.05 | 1.85 | magna |
| fixed | cond | | (Intercept) | -4.74 | 0.80 | -6.53 | -3.40 | longispina |
| fixed | cond | | as.factorlongispina_spring1 | 6.96 | 0.80 | 5.52 | 8.67 | longispina |
| ran_pars | cond | island | sd__(Intercept) | 0.82 | 0.77 | 0.03 | 2.79 | longispina |
| ran_pars | cond | island:pool | sd__(Intercept) | 0.65 | 0.45 | 0.03 | 1.68 | longispina |
| ran_pars | cond | year | sd__(Intercept) | 0.41 | 0.31 | 0.02 | 1.16 | longispina |
| fixed | cond | | (Intercept) | -7.40 | 1.70 | -11.39 | -4.67 | pulex |
| fixed | cond | | as.factorpulex_spring1 | 9.99 | 1.88 | 7.00 | 14.38 | pulex |
| ran_pars | cond | island | sd__(Intercept) | 1.76 | 1.11 | 0.30 | 4.56 | pulex |
| ran_pars | cond | island:pool | sd__(Intercept) | 1.53 | 0.73 | 0.32 | 3.20 | pulex |
| ran_pars | cond | year | sd__(Intercept) | 0.38 | 0.31 | 0.01 | 1.16 | pulex |

#### 2.2 Co-occurrence

Table S2: Estimated coefficients of the model for co-occurrence, where *group* refers to the random effects.

| effect | component | group | term | estimate | std.error | conf.low | conf.high | sample | b |
|---|---|---|---|---|---|---|---|---|---|
| fixed | cond | | (Intercept) | -4.28 | 0.60 | -5.59 | -3.21 | spring | 50.00 |
| fixed | cond | | nr_pools_within | 0.05 | 0.01 | 0.03 | 0.07 | spring | 50.00 |
| fixed | cond | | desiccation_dynamic | 0.00 | 0.21 | -0.44 | 0.40 | spring | 50.00 |
| ran_pars | cond | island | sd__(Intercept) | 1.26 | 0.54 | 0.52 | 2.54 | spring | 50.00 |
| ran_pars | cond | island:pool | sd__(Intercept) | 1.37 | 0.12 | 1.15 | 1.63 | spring | 50.00 |
| ran_pars | cond | year | sd__(Intercept) | 0.39 | 0.12 | 0.18 | 0.64 | spring | 50.00 |
| fixed | cond | | (Intercept) | -5.11 | 0.74 | -6.69 | -3.81 | spring | 100.00 |
| fixed | cond | | nr_pools_within | 0.04 | 0.01 | 0.03 | 0.06 | spring | 100.00 |
| fixed | cond | | desiccation_dynamic | -0.00 | 0.21 | -0.45 | 0.40 | spring | 100.00 |
| ran_pars | cond | island | sd__(Intercept) | 1.50 | 0.57 | 0.70 | 2.88 | spring | 100.00 |
| ran_pars | cond | island:pool | sd__(Intercept) | 1.38 | 0.12 | 1.15 | 1.64 | spring | 100.00 |
| ran_pars | cond | year | sd__(Intercept) | 0.39 | 0.11 | 0.20 | 0.64 | spring | 100.00 |
| fixed | cond | | (Intercept) | -4.28 | 0.63 | -5.65 | -3.16 | summer | 50.00 |
| fixed | cond | | nr_pools_within | 0.04 | 0.01 | 0.02 | 0.06 | summer | 50.00 |
| fixed | cond | | desiccation_dynamic | -0.05 | 0.21 | -0.48 | 0.34 | summer | 50.00 |
| ran_pars | cond | island | sd__(Intercept) | 1.46 | 0.59 | 0.65 | 2.92 | summer | 50.00 |
| ran_pars | cond | island:pool | sd__(Intercept) | 1.24 | 0.11 | 1.03 | 1.48 | summer | 50.00 |
| ran_pars | cond | year | sd__(Intercept) | 0.40 | 0.10 | 0.23 | 0.62 | summer | 50.00 |
| fixed | cond | | (Intercept) | -4.96 | 0.71 | -6.47 | -3.69 | summer | 100.00 |
| fixed | cond | | nr_pools_within | 0.04 | 0.01 | 0.02 | 0.06 | summer | 100.00 |
| fixed | cond | | desiccation_dynamic | -0.05 | 0.21 | -0.48 | 0.34 | summer | 100.00 |
| ran_pars | cond | island | sd__(Intercept) | 1.59 | 0.59 | 0.77 | 3.05 | summer | 100.00 |
| ran_pars | cond | island:pool | sd__(Intercept) | 1.24 | 0.12 | 1.02 | 1.48 | summer | 100.00 |
| ran_pars | cond | year | sd__(Intercept) | 0.40 | 0.10 | 0.23 | 0.63 | summer | 100.00 |

### 3 Supplemental figures

#### 3.1 Linear approximation of the effect of d on local density

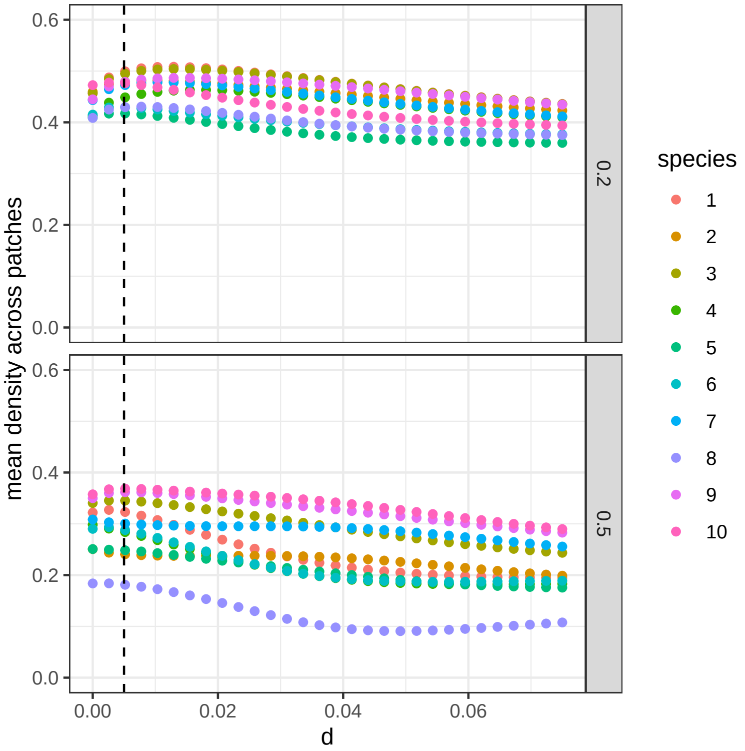

Figure S1: Simulation with the main model for $n = 10$, $p = 50$, two values for $a$ (diffuse competition, identical across patches), and assuming regional equivalence of the growth rate. Dispersal rate $d$ affects mean biomass in a nonlinear way. Only when $d$ is small enough (up to $\approx 0.005$; dashed line) can this effect be approximated by a linear function.

##### 3.2 Preemptive competition

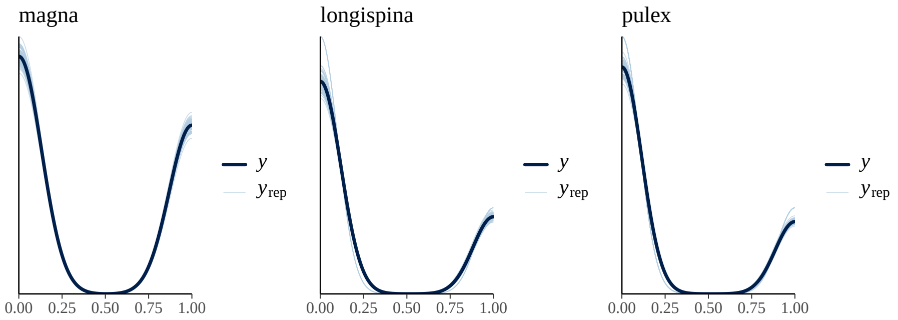

Figure S2: **The model adequately reproduces the observed distribution of occurrences across species.** Posterior predictive checks comparing the observed data ($y$, dark lines) with replicated data from the posterior predictive distribution ($y_{\text{rep}}$, light lines) for each focal species. Close agreement between observed and replicated distributions indicates that the Bernoulli hierarchical model captures the main features of the data, including the prevalence of low probabilities and occasional high-occurrence events.

##### 3.3 Co-occurrence

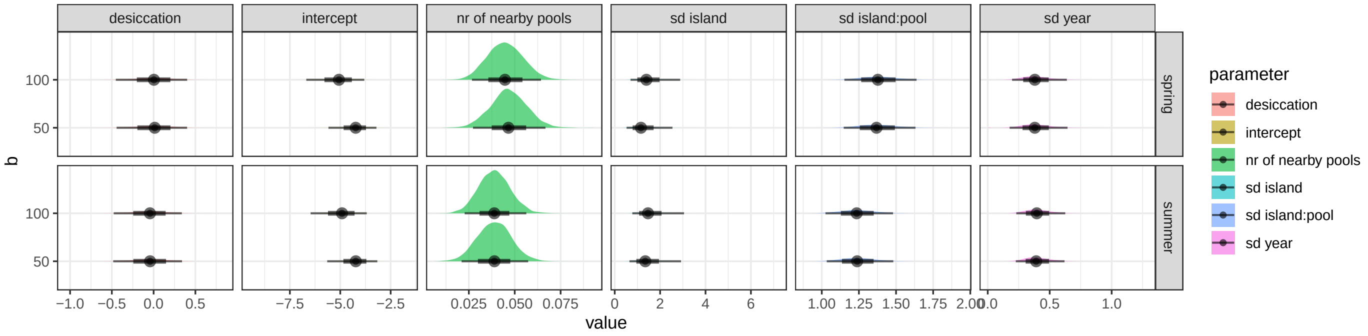

Figure S3: **Occurrence probability increases with local connectivity, while desiccation effects are weak.** Posterior distributions of fixed-effect coefficients and random-effect standard deviations from Bayesian hierarchical models fitted separately for spring and summer and at two spatial scales (50 m and 100 m). The effect of the number of nearby pools is consistently positive across seasons and distances, whereas desiccation effects are centered near zero. Points denote posterior means and thick intervals represent credible intervals; distributions illustrate posterior uncertainty in both fixed and random effects.

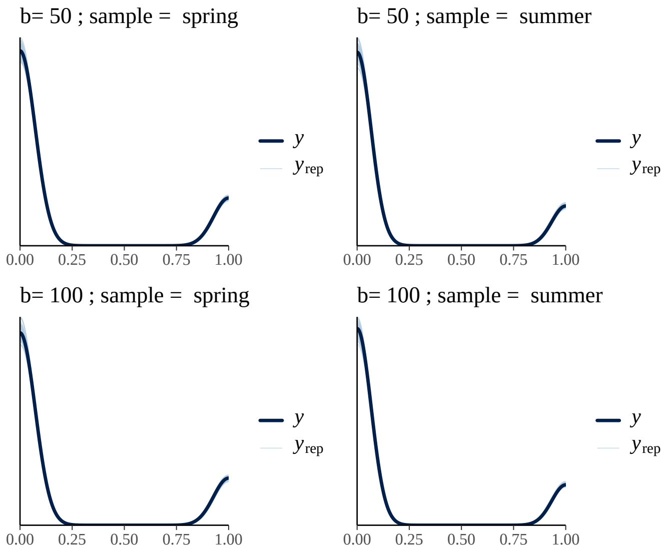

Figure S4: **The proportional-occurrence model reproduces the observed data across seasons and spatial scales.** Posterior predictive checks comparing observed outcomes ($y$, dark lines) with replicated data from the posterior predictive distribution ($y_{\text{rep}}$, light lines) for models fitted at two spatial scales (50 m and 100 m) and in spring and summer. The close correspondence between observed and replicated distributions indicates that the model captures the overall shape and prevalence of low and high occurrence probabilities across conditions.
